# A subset of UPR-induced transmembrane proteins are prematurely degraded during lipid perturbation

**DOI:** 10.1101/178947

**Authors:** Benjamin S.H. Ng, Peter Shyu, Nurulain Ho, Ruijie Chaw, Seah Yi Ling, Guillaume Thibault

## Abstract

**Background:** Phospholipid homeostasis in biological membranes is essential to maintain functions of organelles such as the endoplasmic reticulum. Phospholipid perturbation has been associated to non-alcoholic fatty liver disease, obesity and other metabolic disorders. However, in most cases, the biological significance of lipid disequilibrium remains unclear. Previously, we reported that *Saccharomyces cerevisiae* adapts to lipid disequilibrium by upregulating several protein quality control pathways such as the endoplasmic reticulum-associated degradation (ERAD) pathway and the unfolded protein response (UPR).

**Results:** Surprisingly, we observed certain ER-resident transmembrane proteins (TPs), which form part of the UPR programme, to be destabilised under lipid perturbation (LP). Among these, Sbh1 was prematurely degraded by fatty acid remodelling and membrane stiffening of the ER. Moreover, the protein translocon subunit Sbh1 is targeted for degradation through its transmembrane domain in an unconventional Doa10-dependent manner.

**Conclusion:** Premature removal of key ER-resident TPs might be an underlying cause of chronic ER stress in metabolic disorders.

## BACKGROUND

Phospholipid homeostasis is crucial in the maintenance of various cellular processes and functions. They participate extensively in the formation of biological membranes, which serve to generate distinct intracellular environments into ordered compartments known as organelles for metabolic reactions, storage of biomolecules, signalling, as well as sequestration of metabolites. Existing as various and distinct species, phospholipids are regulated within relatively narrow limits and their composition in biological membranes among organelles differs significantly [1].

Perturbation of the two most abundant phospholipids, phosphatidylcholine (PC) and phosphatidylethanolamine (PE), can lead to various disease outcomes including non-alcoholic fatty liver disease (NAFLD) [2–5], type II diabetes (T2D) [6], as well as cardiac and muscular dystrophies [7]. Being highly abundant in biological membranes, the perturbation of PC and PE levels results in endoplasmic reticulum (ER) stress [8]. For instance, an elevated PC/PE ratio in obesity was found to contribute to the development of NAFLD [9, 10]. Perturbation in phospholipids was shown to cause the premature degradation of the sarco/endoplasmic reticulum Ca^2+^-ATPase (SERCA) ion pump, disrupting calcium homeostasis and resulting in chronic ER stress [9]. This eventually led to hepatic steatosis and liver failure. In another study, mice fed with high fat diet exhibited an increase in gut microbiota enzymatic activity that have been shown to reduce choline [11, 12]. Choline is an essential dietary nutrient primarily metabolised in the liver and used for the synthesis of PC. Similarly, choline deficiency may play an active role in the development of insulin resistance. However, the development of chronic ER stress and metabolic diseases from lipid perturbation (LP) remains largely unknown.

In *Saccharomyces cerevisiae, de novo* synthesis of PC is catalysed by the enzymes Cho2 and Opi3, and similarly carried out by the homologue of Opi3, PEMT, in mammals (Fig. 1a). Cho2 first methylates PE to *N*-monomethyl phosphatidylethanolamine (MMPE), which is further methylated by Opi3 to PC through the intermediate *N,N*-dimethyl phosphatidylethanolamine (DMPE). Alternatively, PC is synthesised from choline, when available, through the Kennedy pathway. Both pathways are highly conserved from yeast to humans. It has been reported that *PEMT^−/−^* mice develop NAFLD within three days of choline deficient diet [5]. Previously, we developed a LP yeast model to mimic NAFLD by deleting the gene *OPI3* [13].

**Figure 1.**
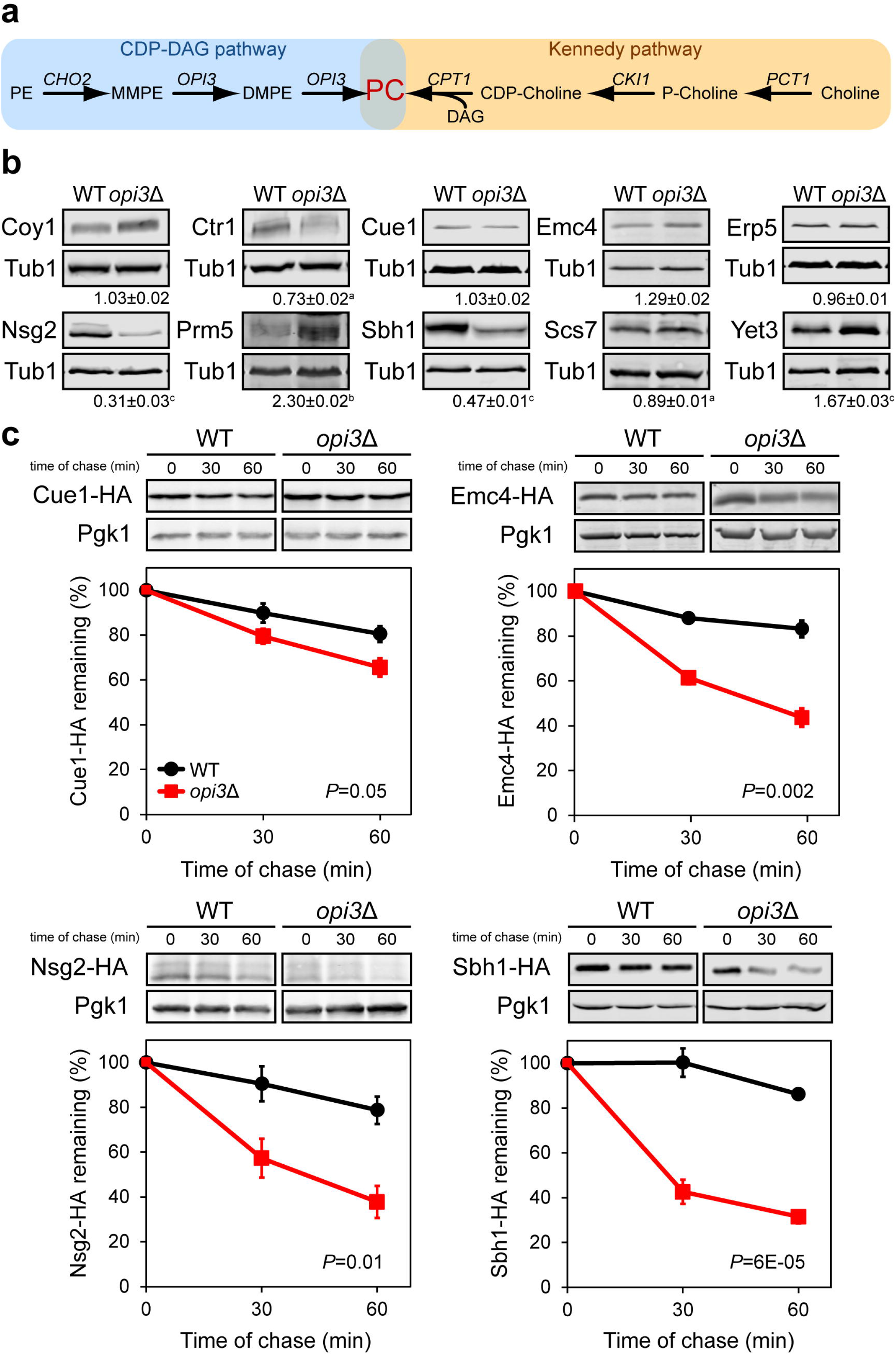
A subset of ER transmembrane proteins is prematurely degraded under lipid imbalance. (**a**) Metabolic pathways for the synthesis of phosphatidylcholine in S. *cerevisiae*. PE, phosphatidylethanolamine; MMPE, N-monomethyl phosphatidylethanolamine; DMPE, N,N-dimethyl phosphatidylethanolamine; PC, phosphatidylcholine; DAG, diacylglycerol; CDP-choline, cytidine diphosphate-choline; P-choline, phosphate-choline. (**b**) Steady state level of transmembrane proteins. Equal cell numbers were harvested. Proteins were separated by SDS-PAGE and detected by immunoblotting with antibodies against the HA tag and Tub1 as loading control. ^a^*P*<0.05, *P*<0.01, ^c^*P*<0.005, Student’s t test. (**c**) Degradation of HA-tagged proteins was analysed after blocking protein translation with cycloheximide. Proteins were separated by SDS-PAGE and detected by immunoblotting with antibodies against the HA tag and Pgk1 as loading control.

The unfolded protein response (UPR) is a stress response pathway monitoring ER stress to restore cellular homeostasis [14]. Upon accumulation of misfolded proteins, the UPR is activated and alleviates stress by reversing severe dysfunctions through the upregulation of nearly 400 target genes in yeast [15]. Major targeted regulatory pathways includes cytosolic protein quality control (CytoQC), ER-associated degradation (ERAD), protein translocation, protein modification and phospholipid biosynthesis [15, 16]. By increasing ER protein folding capacity and enhanced clearance of misfolded proteins coupled with a general attenuation of protein translation [17], the UPR aims to achieve ER homeostasis.

Recently, it was demonstrated that the UPR is essential in alleviating ER stress in lipid dysregulated cells to maintain protein biogenesis, protein quality control and membrane integrity [13, 18–20]. LP, by the absence of *CHO2* or *OPI3*, exhibits synthetic lethality with the sole UPR signalling transducer in yeast, *IRE1*, as well as its downstream transcription factor *HAC1* [19, 20]. LP has been well characterised to induce ER stress [21–23], and the failure of the UPR to restore homeostasis is implicated in several human diseases [24–26]. This clearly establishes the critical role of the UPR in buffering the lethal effects of LP to ensure cell survival.

In this study, we observed certain ER-resident transmembrane proteins (TPs), part of the UPR programme, to be prematurely degraded under LP. First, we demonstrated that LP affects the ER membrane which results in the destabilisation of the TPs. Furthermore, we elucidated the mechanism of how one such TP, Sbh1, gets recognised for degradation through ERAD. Our findings reveal that under LP, Sbh1 transmembrane degron becomes accessible to the Doa10 complex leading to its premature degradation.

## RESULTS

### A subset of transmembrane proteins is destabilised during lipid perturbation

Global transcriptional and proteomic analyses from our previous work indicated a dramatically altered biochemical landscape in yeast cells under LP [13]. Among these, 66 proteins were identified to be transcriptionally upregulated, yet displayed a decrease in protein abundance (Table S1), including 11 ER-resident TPs. From these, we analysed the steady-state levels of ten TP candidates in cells under LP using the PC-deficient strain *opi3*Δ (Fig. 1a-b) [13]. Coy1, Cue1 and Erp5 exhibited similar protein steady-states in *opi3*Δ and WT, while Ctr1, Nsg2, Sbh1 and Scs7 had significantly lower steady-state levels. Surprising, Emc4, Prm5 and Yet3 showed higher steady-state protein levels. To exclude possible cellular functions affected from LP such as transport and secretion, we focused on ER-resident proteins by focusing on Cue1, Emc4, Nsg2, and Sbh1. Cue1 is an essential component of the ERAD pathway [27]. Emc4 is a member of the conserved ER transmembrane complex (EMC) and is required for efficient folding of proteins in the ER [21, 28]. The EMC is also proposed to facilitate the transfer of phosphatidylserine from the ER to mitochondria [29]. Nsg2 regulates the sterol-sensing protein Hmg2 [30]. Lastly, the β subunit of the Sec61 ER translocation complex, Sbh1, is highly conserved in eukaryotes and plays a role in the translocation of proteins into the ER [31–33]. Sbh1 is non-essential for translocation but leads to a defect in this process when deleted in conjunction with its paralogue, Sbh2 [34].

To assess the stability of TP candidates during LP, cycloheximide chase assay was performed in WT and *opi3*Δ. strains. Half-lives of Emc4, Nsg2, and Sbh1 were found to be significantly reduced under lipid disequilibrium (Fig. 1c). No significant decrease in Cue1-HA protein level was detected in *opi3*Δ although the decrease was reproducible. One hour after attenuating protein translation, levels of Emc4, Nsg2, and Sbh1 were found to be 27%, 41%, and 58% lower in *opi3*Δ., respectively, compared to WT. This suggests that the UPR programme transcriptionally upregulates genes to restore ER homeostasis under LP, while a subset of TPs is recognised and targeted for degradation.

### A subset of ER-localised transmembrane proteins is destabilised by a decrease in phosphatidylcholine

To ensure that Cue1, Emc4, Nsg2, and Sbh1 remain as integral ER membrane proteins during lipid perturbation, we verified their localisation at the ER (Fig. 2a) and their insertion into cellular membranes (Fig. 2b) in *opi3*Δ. cells. Together, these results suggest that integration into the ER membrane is unaffected by PC depletion. To study the topology of these four proteins, we performed proteinase K (PK) digestion from isolated microsomes (Fig. 2c). In WT cells, the C-termini HA tags of Cue1-HA, Emc4-HA and Nsg2-HA are oriented towards the cytosol. Thus, the HA tag will be cleaved off if the proper topology is preserved, while the detection of a HA-bearing peptide after PK digestion indicates an inverted topology. The three proteins were found to be fully digested under LP and the predicted smaller protein fragments of 23.7, 8.53, and 5.8 kDa were not detected for Cue1-HA, Emc4-HA, and Nsg2-HA, respectively, in both WT and *opi3*Δ. Sbh1-HA is a tail-anchored protein where the C-termini HA tag is found in the ER lumen. The predicted protein fragment of 10.5 kDa after PK digestion was detected in both WT and *opi3*Δ. strains, indicative of its correct membrane topology. Typically, tail-anchored proteins are tagged at the N-termini as the C-termini interacts with the Get complex for insertion into the ER membrane [35]. This result shows that, along with alkaline carbonate extraction (Fig. 2b), adding a C-terminus HA tag to Sbh1 does not interfere with its integration into the ER membrane. The four TPs were fully digested in the presence of the non-ionic detergent Nonidet P-40 (NP40). Together, these findings suggest that the four TPs are prematurely targeted for degradation once they are fully translated and integrated into the ER membrane.

**Figure 2.**
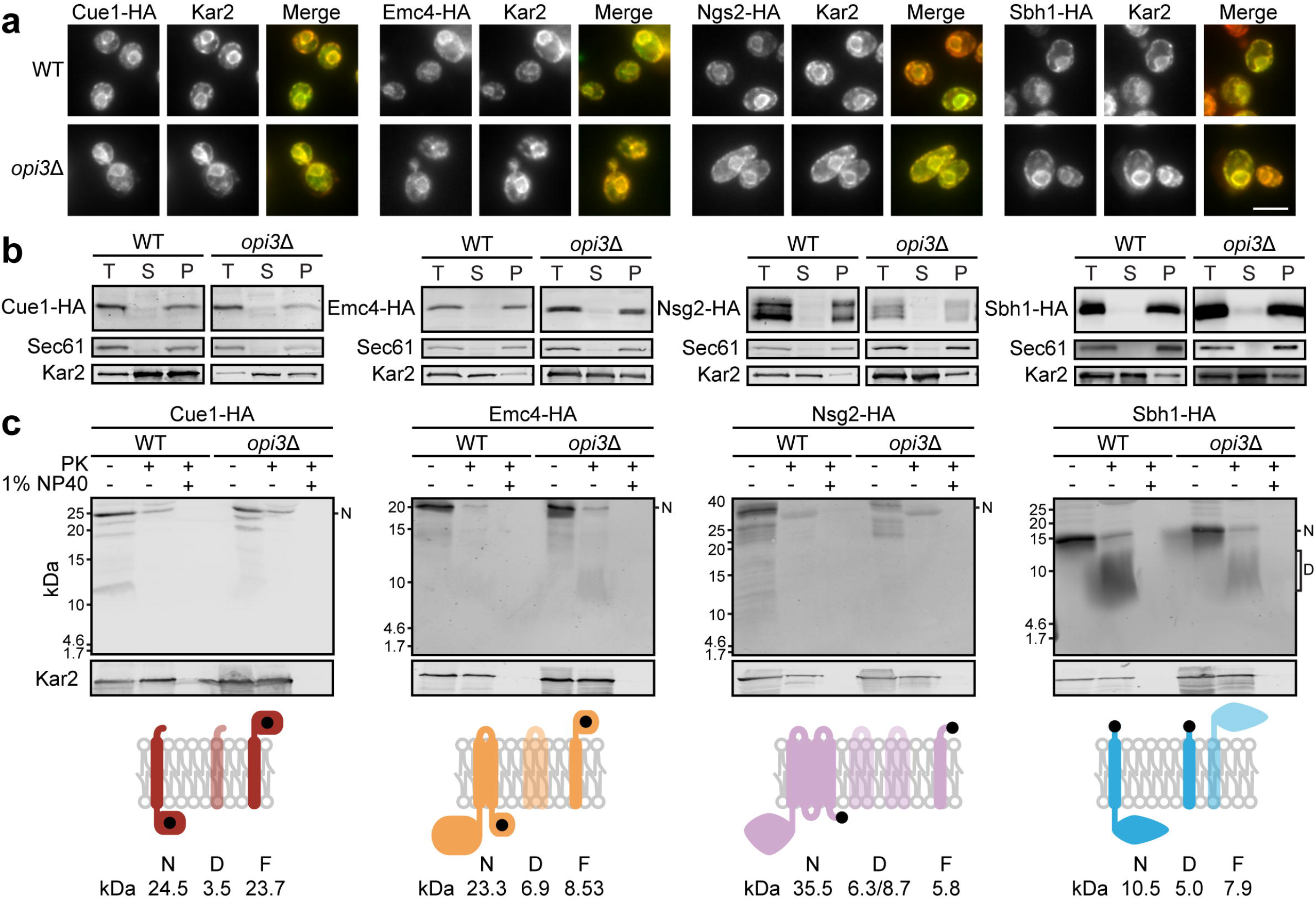
Transmembrane proteins are destabilised by the decrease in phosphatidylcholine synthesis. (**a**) Protein candidates were detected using antibodies against HA tag and Kar2 as ER marker. Scale bar, 5 μm. (**b**) Membranes prepared from wild type and *opi3*Δ cells expressing HA-tagged proteins were treated with 0.1 M sodium carbonate, pH 11, for 30 min on ice. A portion was kept as the total fraction (**T**), and the remaining was subjected to centrifugation at 100,000 X g. Supernatant (S) and membrane pellet (P) fractions were collected and analysed by immunoblotting. Proteins were detected using anti-HA antibody. Kar2 and Sec61 serve as soluble and integral membrane protein controls, respectively. (c) Membranes prepared from WT and *opi3*Δ cells expressing HA-tagged proteins were treated with 1 mg/ml proteinase K, for 30 min at 37°C, with or without 1% NP40. HA-tagged proteins were precipitated with 10% TCA, separated by SDS-PAGE and detected by immunoblotting with HA antibody. Expected protein molecular weights are shown below for nondigested (N), digested (D), and flipped and digested (F). The orientation of the HA tag is shown as black dot. Fragments missing the HA tag and are therefore undetectable are illustrated with transparency. The ER lumen and cytosol are at the top and bottom of the membrane, respectively.

To further confirm the four TPs are destabilised specifically from low PC levels, their degradation was monitored in cells grown in the presence of choline to restore PC homeostasis (Fig. 1a) [13, 36]. Choline supplementation significantly stabilised Cue1-HA, Emc4-HA, Nsg2-HA, and Sbh1-HA in *opi3*Δ to the levels of WT (Fig. 3a). Subsequently, we concentrated our effort on Sbh1 to better understand how it is targeted for premature degradation during LP.

**Figure 3.**
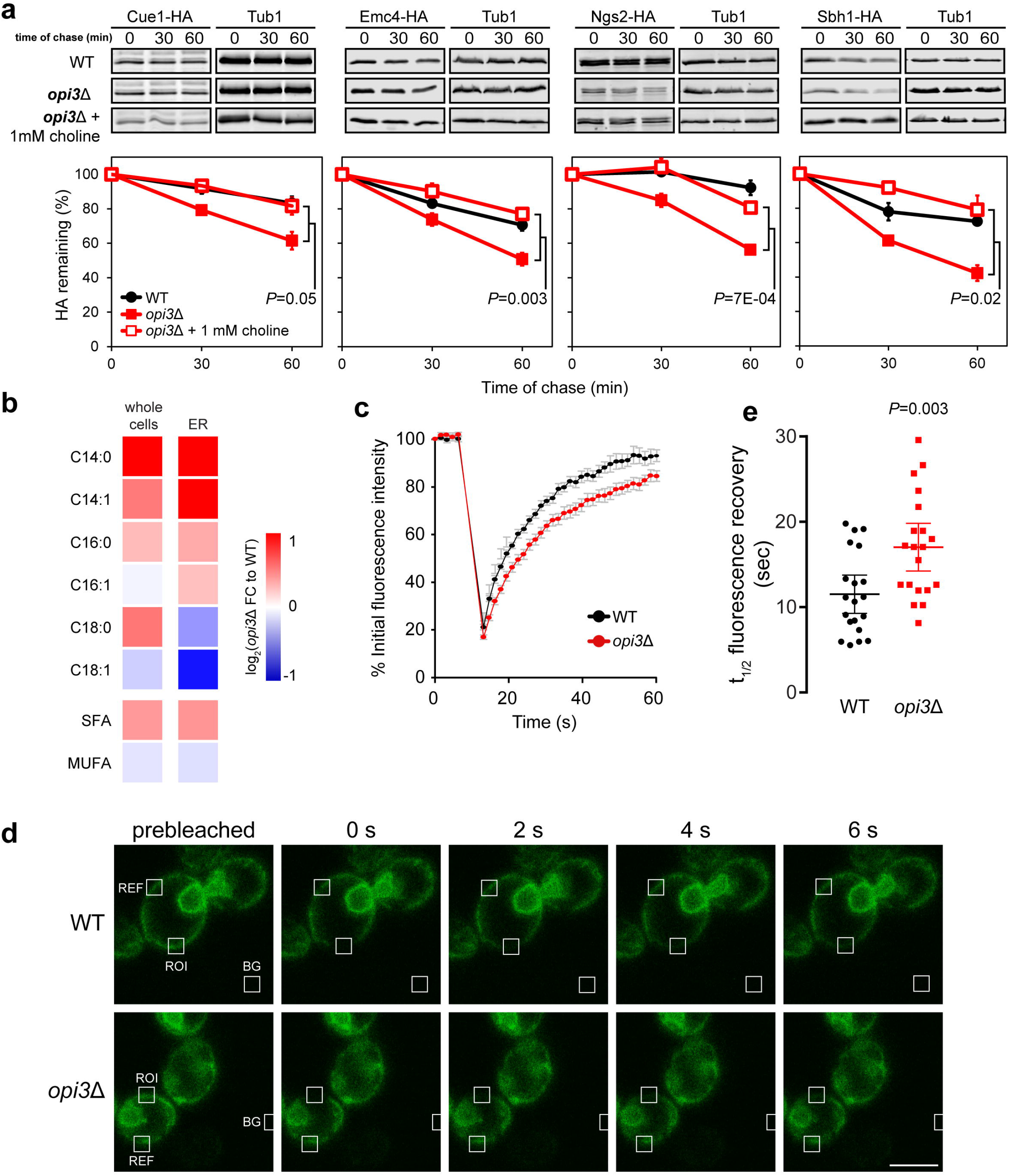
Sbh1 is destabilised from increased membrane fluidity of the ER membrane. (**a**) Cell were grown with or without 1 mM choline before addition of cycloheximide. Time points were taken as indicated. Proteins were separated by SDS-PAGE and detected by immunoblotting with antibodies against the HA tag and Tub1 as loading control. (**b**) Heat map of log_2_-transformed fold changes (FC) in fatty acids (FA) in *opi3*Δ as compared to WT. FAs in whole cells and microsomes (ER) of WT and *opi3*Δ were quantified by gas chromatography after FAME derivatisation. (**c-e**) Fluorescence recovery after photobleaching using Sec63-sGFP in WT and *opi3*Δ. (**c**) Averages of Sec63-sGFP signal intensity from 20 cells are plotted over a 60-second period. (**d**) Fluorescence intensity was monitored from the white boxes ROI (region of interest), REF (reference), and BG (background). Scale bar, 5 μm. A region of the cortical ER of live cells were photobleached and recovery points at 1.57 s intervals were taken. (**e**) The time elapsed for the half-maximal fluorescence recovery (t_½_) was calculated and plotted. Student’s t test compared to WT.

The UPR is strongly activated in response to LP [13, 21]. In *opi3*Δ, the UPR activation is constitutively elevated and unresolved, thereby referred to as chronic ER stress [13, 37]. To ensure that Sbh1 is not destabilised as a consequence of strong UPR activation, we introduced a constitutively active form of the downstream effector, *HAC1^i^*, into WT cells [16, 38]. As expected, *HAC1^i^*-induced UPR activation did not further destabilise Sbh1 in WT cells (Additional file 1: Fig. S1a). Noticeably, steady-state Sbh1 protein level is higher in UPR-activated WT cells as *SBH1* is upregulated from the UPR programme [13, 15]. Additionally, yeast cells can mount an intact UPR in the absence of *SBH1* (Additional file 1: Fig. S1b). Thus, this indicates that the UPR programme in *opi3*Δ is not sufficient to drive premature Sbh1 degradation.

### Changes in ER membrane fluidity is sufficient to destabilise Sbh1

To narrow down the specific effect of LP that might contribute to the premature degradation of Sbh1, we analysed the fatty acid (FA) composition of whole cells and fractionated microsomes. Overall, there was a general increase of cellular and microsomal (ER) saturated fatty acids (SFAs) and decrease of monosaturated fatty acids (MUFAs) in *opi3*Δ. when compared to WT (Fig. 3b). A significant decrease of oleic acid (C18:1) was also observed in *opi3*Δ. microsome fraction compared to that of WT. In addition to FA remodelling, the intermediate for the synthesis of PC from PE, MMPE, largely accumulates with the deletion of *OPI3* as we previously reported (Fig. 1a) [13]. A large MMPE increase is expected to induce negative membrane curvature stress as has been reported for PE [39]. The remodelling of FA saturation state could be another adaptive response to alleviate membrane curvature stress in *opi3*Δ [40, 41], as FA saturation states of biological membranes are highly linked to membrane fluidity [42–44].

To better understand the impact of membrane remodelling on TPs behaviour, we monitored the dynamics of the ER-resident membrane protein Sec63-sGFP by fluorescence recovery after photobleaching (FRAP) [45]. A region of the cortical ER is photobleached and signal recovery correlates with Sec63-sGFP mobility. The recovery of Sec63-sGFP fluorescence was significantly slower in *opi3*Δ compared to WT suggesting rigidity of the ER membrane (Fig. 3c-e). This result is consistent with previous reports on the effect of decreased PC/PE ratio in stiffening membranes [40, 46]. Taken together, it suggests that a decrease in membrane fluidity might prevent TPs to associate to their interacting partners following translation and resulting in premature degradation.

### Sbh1 binding to interacting partners is compromised under lipid imbalance

To further characterise the effect of LP on Sbh1 stability, we performed the split-ubiquitin based membrane yeast two hybrid (MYTH) screen in WT and *opi3*Δ cells to identify changes in Sbh1 membrane protein interactome [47, 48]. The reporter moiety was added at the N-terminus of Sbh1 (TF-C_ub_-Sbh1) and did not compromise its ER localisation (Additional file 1: Fig. S2a). Strains expressing the Sbh1 bait were transformed with a yeast prey genomic plasmid library in which open reading frames are fused to sequences encoding the cognate reporter moiety [49]. A total of 49 and 14 putative Sbh1-interacting proteins were identified in WT and *opi3*Δ, respectively (Additional file 1: Fig. S2b). To eliminate false positive interactors, a bait dependency test was done using the singlepass transmembrane domain of the human T-cell surface glycoprotein CD4 tagged to C_ub_-LexA-VP16 [49]. In WT, we identified 38 bona fide Sbh1 interactors including previously reported interactors Ost4, Sec61, Spc2, Ssb1, Sss1, and Yop1 (Fig. 4a) [50–53]. Sbh1 was also found to interact with membrane proteins involved in sterol biogenesis (Erg4, Erg24 and Nsg1) and fatty acid elongation (Elo2 and Tsc13). On the other hand, only 13 proteins were found to interact with Sbh1 in *opi3*Δ cells (Fig. 4b). No interaction of Sbh1 with Sec61 and Sss1 was detected in *opi3*Δ. This suggests that Sbh1 could be dissociated from the Sec61 complex under LP, and therefore causes its premature degradation. This is consistent with the finding that Sbh2, the paralogue of Sbh1, becomes destabilised and degraded rapidly when unbound to the Sec61-like complex Ssh1 [54]. Similarly, Sbh1 was found to interact with proteins of the ERAD pathway under LP (Fig. 4b). Sbh1 interactors include the membrane-embedded ubiquitin-protein ligase Doa10 which is part of the ERAD Doa10 complex [55, 56]. As the Doa10 complex is generally specific for substrates containing cytosolic lesions (ERAD-C) [57], it suggests that a polypeptide stretch of Sbh1 might become exposed on its cytosolic side under LP making it susceptible to ubiquitination. Subsequently, targeted substrates for degradation are polyubiquitylated in the cytosol by the addition of Lys-11-linked ubiquitin (Ubi4), a protein identified to interact with Sbh1 exclusively in *opi3*Δ cells. The AAA^+^ ATPase protein Cdc48 was also found to interact with Sbh1 in *opi3*Δ cells (Fig. 4b). Ubiquitylated substrates are retro-translocated to the cytosol by the action of the Cdc48 complex and targeted to the proteasome for degradation [58, 59]. Another important player of the ERAD pathway, Png1, was found to exclusively interact with Sbh1 under LP. Png1 catalyses the deglycosylation of misfolded glycoproteins, and is a critical step for ERAD substrates modification to be fit for proteasomal degradation [60]. Together, the MYTH screening results suggest that a change in membrane properties lead to the dissociation of Sbh1 from the Sec61 complex, resulting in its rapid degradation through the ERAD-C complex.

**Figure 4.**
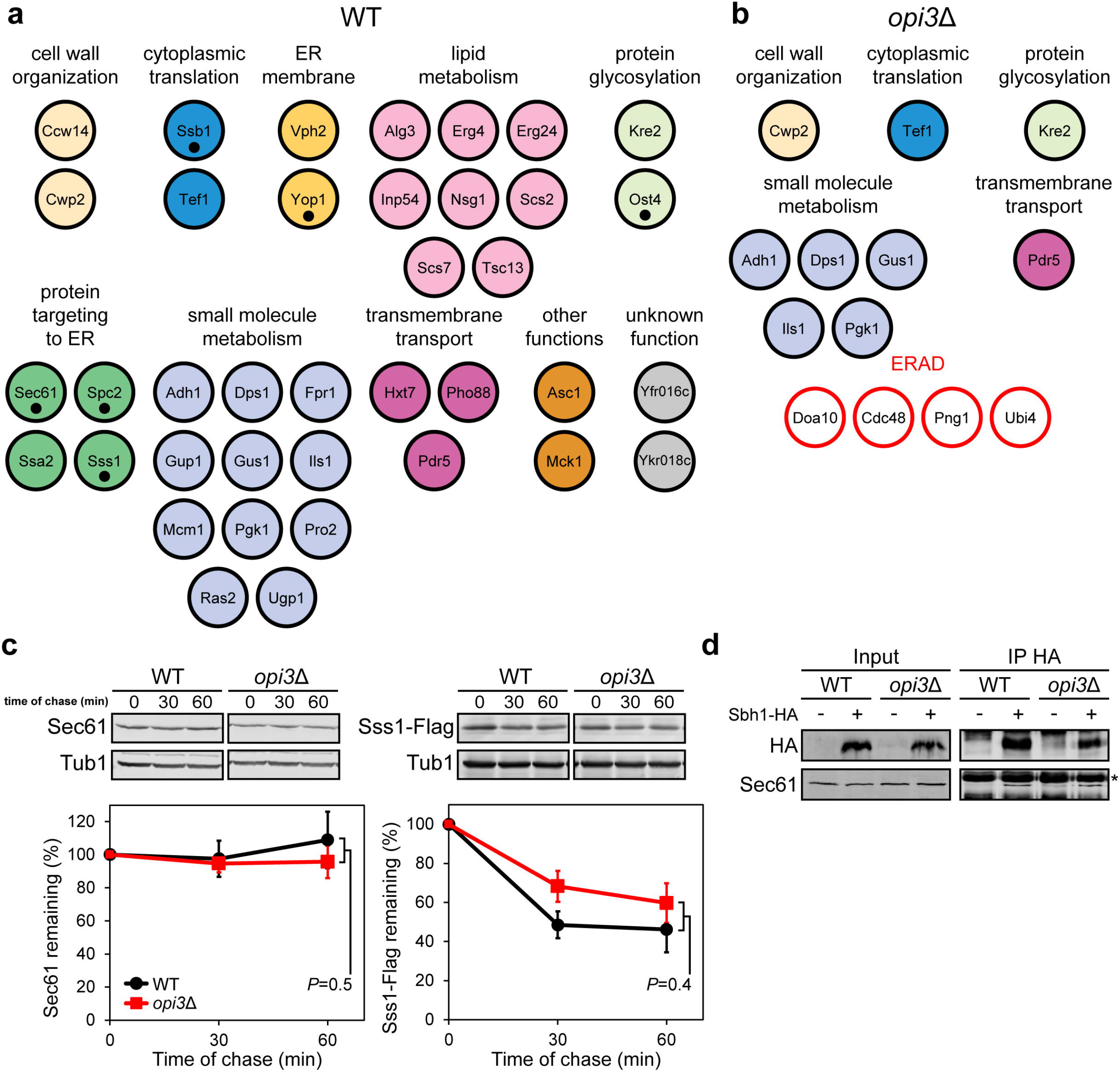
Sbh1 binding to interacting parters is compromised under lipid imbalance. (**a,b**) Proteins identified as interacting partners of N-termini reporter tagged Sbh1 (TF-C_ub_-Sbh1) by the MYTH method in WT (A) and *opi3*Δ (B) cells. ERAD factors were only detected in *opi3*Δ and are denoted in red. Previously reported interactors of Sbh1 are indicated with black dots. (c) The degradation of Sec61 or Sss1-Flag was analysed in WT and *opi3*Δ cells after blocking translation with cycloheximide. Proteins were separated by SDS-PAGE and detected by immunoblotting with antibodies against Sec61 or Flag tag and Tub1 as loading control. (**d**) Immunoprecipitation of Sbh1-HA with protein G beads were analysed in WT and *opi3* native cell lysates. Eluted and input fractions were resolved by SDS-PAGE, transferred to nitrocellulose membrane, and analysed by immunoblotting with antibodies against Sec61 and the HA tag after the release of HA bound Sbh1 with HA peptide.

To ensure levels of the Sec61 complex subunits other from Sbh1 remain unchanged under lipid perturbation, we carried out cycloheximide chase assay to follow the stability of Sec61 and Sss1-Flag. Both Sec61 and Sss1 were found to be stable in *opi3*Δ as in WT in agreement with our previous proteomic data (Fig. 4c) [13]. To assess the interaction of Sbh1 with Sec61 complex on the ER membrane under LP, native co-immunoprecipitation (co-IP) was performed (Fig. 4d). In contradiction to the MYTH screen results, Sec61 was found to interact stably with Sbh1-HA in both WT and *opi3*Δ strains. The discrepancy could be due to the difference in membrane dynamics *in vivo* and *in vitro* from the MYTH and co-IP assay, respectively.

### Sbh1 is destabilised from its transmembrane domain and degraded in a Doa10-dependent manner

To validate that Sbh1 is degraded in a Doa10-dependent manner, we carried out cycloheximide chase assay to monitor Sbh1 stability in different ERAD mutants. Sbh1 was found to be fully stabilised in *opi3Δdoa10Δ* but not in *opi3*Δ*hrd*1Δ and *opi3*Δ*usa*1Δ mutants (Fig. 5a). Hrd1 and Usa1 are both part of the Hrd1 complex which recognises lesions within the luminal domains of membrane and soluble proteins (ERAD-L) and those found within transmembrane region (ERAD-M) [61]. As some misfolded proteins in the ER are routed to the vacuole for degradation, we confirmed that Sbh1 degradation under LP is independent of the vacuolar pathway as shown by a similar degradation profile in *opi3*Δ*pep4*Δ. Conversely, Sbh1 degradation showed dependency on Cue1, a conserved element in both the Doa10 and Hrd1 complexes (Additional file 1: Fig. S3). Together with the MYTH data, it suggests that Sbh1 is exclusively targeted for degradation by the ERAD Doa10 complex.

**Figure 5.**
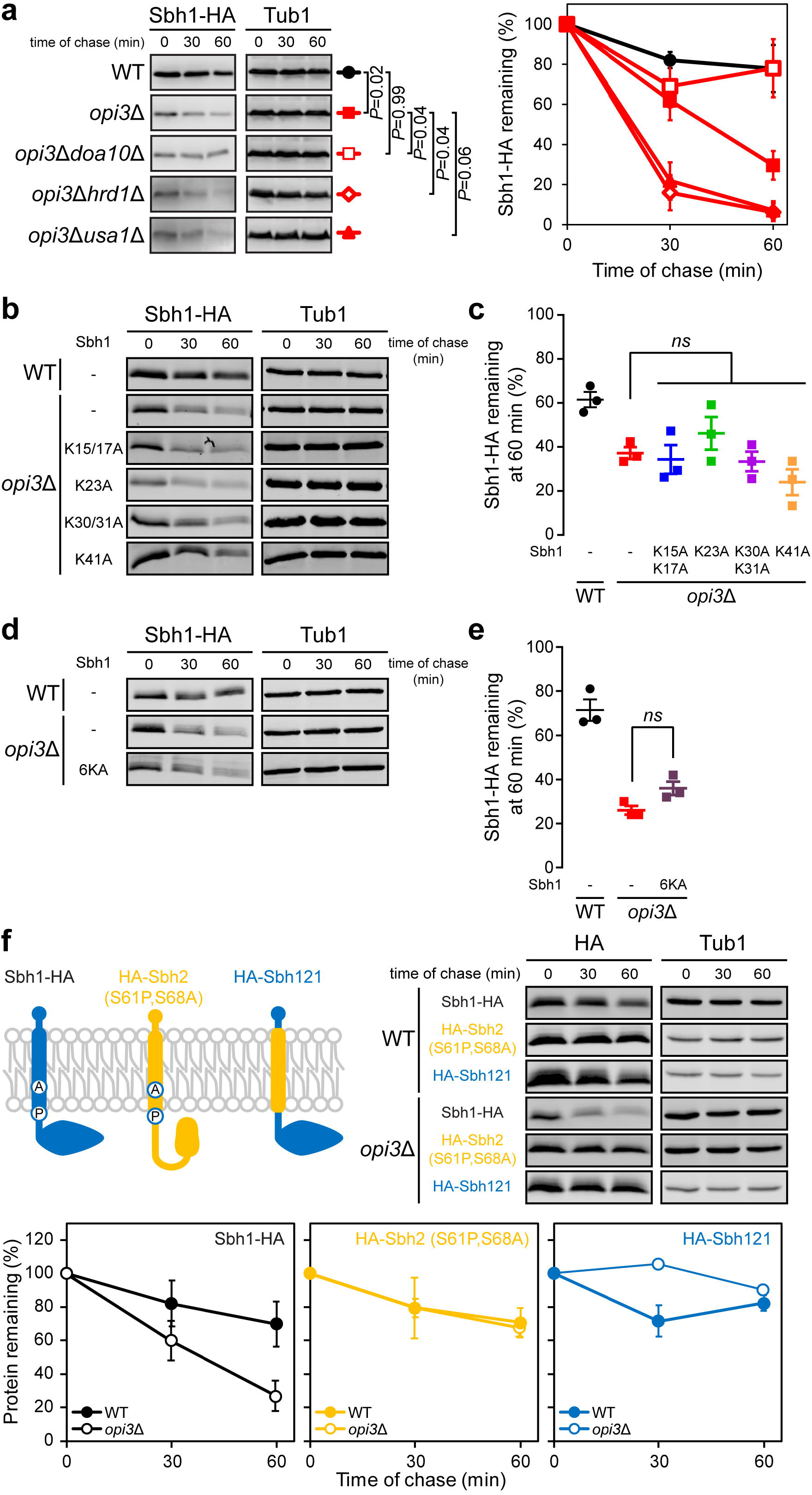
Sbh1 is destabilised from its transmembrane domain and degraded in a Doa10-dependent manner. (**a**) The degradation of Sbh1-HA was analysed in WT, *opi3*Δ, *opi3*Δ*doa10*Δ, *opi3*Δ*hrd1*Δ, and *opi3*Δ*usa1*Δ cells after blocking translation with cycloheximide. Proteins were separated by SDS-PAGE and detected by immunoblotting with antibodies against the HA tag and Tub1 as loading control. (**b**) The degradation of Sbh1-HA in WT and *opi3*Δ cells or Sbh1 cytosolic lysine mutant in *opi3* cells treated as in a. (**c**) Sbh1 percentage remaining at the 60 min time point from b. (**d**) The degradation of Sbh1-HA in WT and *opi3* cells or Sbh1 all cytosolic lysine mutated to alanine [Sbh1(6KA)] in *opi3*Δ cells treated as in a. (**e**) Sbh1 percentage remaining at the 60 min time point from d. (**f**) The degradation of mutant Sbh2 with amino acids 61 and 68 mutated to proline and alanine, respectively [HA-Sbh2(S61P,S68A)], and chimeric Sbh1 protein with its transmembrane domain replaced with that of Sbh2 (HA-Sbh121) in WT and *opi3*Δ cells treated as in a. The ER lumen and cytosol are at the top and bottom of the membrane, respectively.

To further elucidate how Sbh1 might be targeted for degradation by the Doa10 complex during LP, we mutated Sbh1 cytosolic lysine residues to alanine separately [Sbh1(K15A,K17A), Sbh1(K23A), Sbh1(K30A,K31A), and Sbh1(K41A)] and combined [Sbh1(6KA)]. The E3 ubiquitin-protein ligase Doa10 has been extensively reported to recognise ER proteins with cytosolic lesions resulting in the transfer of ubiquitin to lysine residues [62–69]. The degradation rates of Sbh1(K15A,K17A), Sbh1(K23A), Sbh1(K30A,K31A), and Sbh1(K41A) expressed in *opi3*Δ cells were similar to unmutated Sbh1 (Fig. 5b,c). Similarly, Sbh1(6KA) destabilisation was comparable to unmutated Sbh1 in *opi3*Δ strain (Fig. 5d,e). Together, these findings suggest that Sbh1 is targeted for degradation by the Doa10 complex independently from the ubiquitination of its cytosolic domain. Yeast paralogue of Sbh1, Sbh2, is degraded by Doa10 through an intramembrane degron [54]. Thus, we examined the degradation of Sbh1 containing the transmembrane domain of Sbh2 in *opi3* strain (Fig. 5f). Replacing the transmembrane domain of Sbh1 was sufficient to stabilise it during LP suggesting the degron recognised by Doa10 is within the lipid-embedded Sbh1 α-helix [54]. To further validate this finding, we used a stable Sbh2 mutant wherein the two non-conserved amino acids of the transmembrane domain of Sbh2 have been mutated from serine to proline and alanine at positions 61 and 68, respectively [Sbh2(S61P,S68A)]. These two point mutations drive Sbh2 native interaction from the Ssh1 translocon to the Sec61 translocon. As previously reported, Sbh2(S61P,S68A) was stable in WT cells (Fig. 5f). Unexpectedly, Sbh2(S61P,S68A) was similarly stable in *opi3*Δ cells, suggesting the Sec61 translocon maintains its ability to interact with the non-essential β subunit. Together these findings suggest that the Doa10 complex recognises the Sbh1 transmembrane degron that becomes accessible during LP perhaps due to the change in the ER membrane composition.

## DISCUSISON

The strong association between obesity and non-alcoholic fatty liver disease (NAFLD) in human populations is evident of the importance of lipid regulation in determining the emergence of fatty liver pathogenesis: NAFLD is now the most common cause of chronic liver enzyme elevation and cryptogenic cirrhosis, as a result of increased obesity [70, 71]. Total PC is consistently decreased in NAFLD and non-alcoholic steatohepatitis (NASH) liver samples from human patients and mouse models [8, 72, 73], and it correlates with a decrease of the enzyme required for *de novo* synthesis of PC in the liver, PEMT [9, 73]. Concurrently, chronic ER stress and the activation of the UPR are both associated with NAFLD pathologies [9, 74, 75]. Despite these connections, little is known on the effect of phospholipid perturbation on pathways of the ER. Thus, we sought to better understand how the ER fails to reach homeostasis under chronic PC depletion and how the protein quality control machinery is implicated using our previously reported yeast model system [13].

The proteostasis network undergoes extensive remodelling upon PC depletion in yeast [13]. Although a large subset of proteins is increased in these stressed cells, we noticed that key proteins are rapidly degraded and are indeed sensitive to phospholipid variations. Out of the 66 proteins which displayed decreased protein abundance despite being genetically upregulated, 40% are transmembrane proteins (TPs). As 30% of the proteome is predicted to be integral or peripheral membrane proteins [52], it suggests that TPs are more sensitive to LP compared to other types of proteins. Among the identified TPs, a large proportion are ER-resident proteins suggesting this organelle is more vulnerable to the effects of LP, and that this in turn affects TP integrity in the ER. The virtual absence of sterol at the ER, a key regulator of membrane fluidity, might contribute to its susceptibility to change in the biophysical properties of the membrane through lipid variation {Zinser, 1993 #871 ?Weete et al., 2010;Subczynski, 2017 #943}.

We sought to investigate changes in membrane properties under LP that caused the destabilisation of a subset of TPs. PC is cylindrically shaped with a cross-sectional area for the head-group similar to its constituent acyl chain tails, generating minimal curvature and forming flat lamellar phase phospholipid bilayers [76]. PE is classified as cone-shaped lipid forming non-lamellar membrane structure as it generates negative membrane curvature [39]. The phospholipid intermediate MMPE becomes highly abundant under the ablation of *OPI3*, and being mono-methylated, it has physical properties more similar to PE (Fig. 1a). The increase in membrane curvature from the replacement of PC to MMPE may induce cells to decrease their FA chain lengths in accordance to the seminal Helfrich theory of membrane bending elasticity (Fig. 3b) [41]. A more pronounced remodelling of the FA chain length in the ER over whole cell suggests either the ER is more susceptible to LP due to the minimal presence of ergosterol at the ER [77] or cells respond more aggressively to the ER membrane bilayer disruption to alleviate ER stress. Accordingly, a rise in membrane lipid packing from elevated saturated fatty acids will reduce the propensity to form curvatures.

However, the remodelling of the ER to alleviate negative membrane curvature stress, induced from high PE and MMPE levels, can impose further challenges to cells. An elevation in saturated fatty acid chains decreases ER membrane fluidity (Fig. 3b) [78] which might be partially due to the absence of the rich unsaturated fatty acid provider, PC [79, 80]. Additionally, the replacement of PC with MMPE contributes to the stiffening of the membrane [46]. Thus, these changes combined with the relatively low abundance of ergosterol at the ER membrane bilayer make this organelle particularly susceptible to PC level variations. Indeed, this change in the ER membrane led to the premature degradation of Sbh1 by the Doa10 complex through a degron within the transmembrane domain of Sbh1. The loss of Sbh1 interacting partners, during LP, might contribute to its degradation as reported for its yeast paralogue Sbh2 [54]. Dissociation of Sbh2 from the Ssh1 complex (yeast Sec61 paralogue) was proposed to sufficiently drive its Doa10-mediated degradation. Interestingly, none of the Sbh1 cytosolic lysine residues are required for its degradation through the Doa10 complex suggesting Sbh1 might by atypically ubiquitylated as has been reported for the Doa10 substrate Asi2 [81].

Alteration of lipid raft composition at the plasma membrane can lead to loss of protein function and rapid degradation [79, 82, 83]. The rigidity of the ER membrane, from depleting PC, may interfere with Sbh1 conformational changes necessary for its interaction with the Sec61 complex and thus result in its degradation [84]. Alternatively, the stiffening of the lipid membrane may reduce Sbh1 diffusion through the lipid bilayer leading to sustained interaction with the Doa10 complex (Fig. 4 and 6) [54, 85]. Thus, a decrease in PC clearly targets Sbh1 for degradation from a change in the biophysical property of the membrane. It remains to be determined if the LP-induced degradation mechanism of Sbh1 applies to the other destabilized TPs that have been identified (Fig. 1b). Additionally, the absence of PC with its large head-group and the abnormally high presence of PE and MMPE with smaller head-groups at the lipid membrane-cytosol interface should result in Doa10 accessibility of the Sbh1 α-helix degron [54].

The coordinated upregulation of the proteostasis network by the UPR serves as an important stress recovery mechanism that helps cells cope with the otherwise lethal effects of LP [13]. Despite this robust stress response under LP, the UPR programme fails to increase the expression level of a subset of TPs. The premature degradation of these TPs can prevent an effective proteostatic response especially under prolonged LP (Fig. 6). ER stress induced from a temporary lipid perturbation will result in the upregulation of UPR target genes and consequently ER homeostasis. However, in the context of fatty liver, prolonged LP might prevent cells from reaching ER homeostasis by the premature degradation of key UPR target TPs. Therefore, this will lead to chronic ER stress which might contribute to the progression of NAFLD. In addition, the prolonged upregulation of lipogenic transcription factors from the UPR programme may also contribute to liver progression into hepatosteatosis [86].

**Figure 6.**
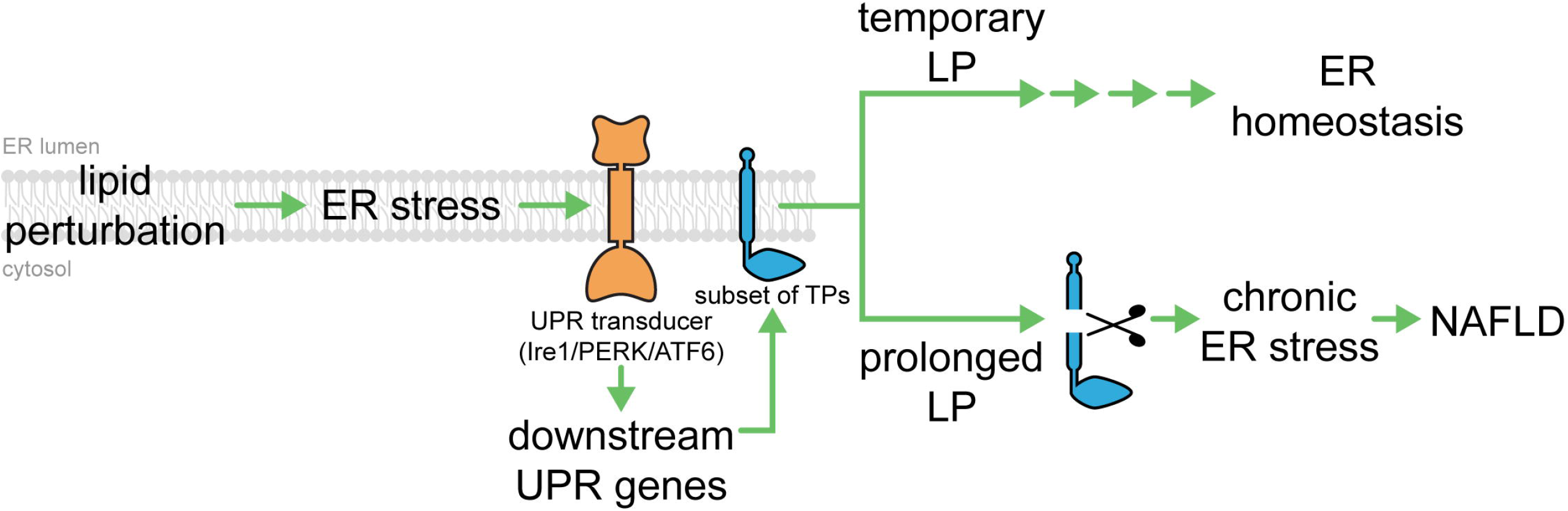
Premature degradation of TPs leads to chronic ER stress and development of NAFLD. Normally, ER homeostasis can be reached from lipid perturbation through the regulation of downstream UPR target genes. UPR transactivator (yellow protein representing Ire1, PERK, or ATF6) senses ER stress from the accumulation of misfolded proteins and/or lipid perturbation. However, under prolonged LP, ER homeostasis could not be achieved due to the premature degradation of a subset of misfolded proteins (blue protein) leading to chronic ER stress, cell death, and eventually the development of NAFLD.

In contrast, disrupting phospholipid homeostasis may be exploited to target pathogens. An increase in phospholipid synthesis is essential for replication of the parasite *Plasmodium falciparum* during the erythrocytic stage [87]. Phospholipid content of parasite-infected erythrocytes dramatically increases during maturation with 85% of newly synthesised phospholipids being PC and PE for growth and cell division [88]. Hence, the inhibition of phospholipid synthesis might be an effective strategy for antimalarial drugs [87, 89]. In addition, *P. falciparum* resistance to artemisinin-based combination therapies (ACTs) is associated to ER stress where the UPR mitigates artemisinin-induced protein damage [90]. Thus, targeting phospholipid biosynthesis in combination with artemisinin might be an efficient strategy to overcome resistance by preventing effective UPR activation in *P. falciparum*. [91]. Similarly, it may be applied to therapeutic strategies against diseases such as cancer where UPR activation is a potent driver of cell division [24, 92].

## CONCLUSIONS

Here, we report that a subset of transmembrane proteins, part of the UPR programme, are prematurely degraded under LP. ER-resident proteins Cue1, Emc4, Nsg2, and Sbh1 topology and integration into the ER are not affected by LP while they are prematurely degraded. By further investigating the β subunit of Sec61 ER translocation complex, Sbh1, we proposed that it is prematurely degraded by the Doa10 complex through the recognition of a specific transmembrane degron. The proper association of Sbh1 with its interacting partners as well as the maintenance of membrane lipid PC level should be sufficient to prevent the Sbh1 degron from being recognised by the Doa10 complex during lipid equilibrium. However, the drastic decrease of PC associated with fatty liver promotes the dissociation of Sbh1 from its interacting partners as well as the exposure of Sbh1 proline 54 leading to its premature degradation in a Doa10-dependent manner. Thus, the premature degradation of a subset of ER-resident TPs during prolonged lipid perturbation might contribute to chronic ER stress associated with NAFLD and NASH.

## METHODS

### Statistics

Error bars indicate standard error of the mean (SEM), calculated from at least three biological replicates, unless otherwise indicated. *P* values were calculated using two-tailed Student’s t test, unless otherwise indicated, and reported as P=value in figures.

### Strains and antibodies

*Saccharomyces cerevisiae* strains used in this study are listed in Additional file 1: Table S2. Strains were generated using standard cloning protocols. Anti-Kar2 polyclonal rabbit antibody and anti-Sec61 polyclonal rabbit antibody were gifts from Davis Ng (Temasek Life Sciences Laboratories, Singapore). Anti-HA mouse monoclonal antibody HA.11 (Covance, Princeton, NJ), anti-Pgk1 mouse monoclonal antibody (Invitrogen), anti-GFP mouse monoclonal antibody (Sigma-Aldrich, St. Louis, MO) anti-tubulin mouse monoclonal antibody 12G10 (DHSB) and anti-LexA polyclonal rabbit antibody (Abcam, Cambridge, United Kingdom) were commercially purchased. Secondary antibodies goat anti-mouse IgG-DyLight 488 (Thermo Fisher, Waltham, MA), goat anti-rabbit IgG-DyLight 550 (Thermo Fisher, Waltham, MA), goat anti-mouse IgG-HRP (Santa Cruz Biotechnology, Dallas, TX), goat anti-rabbit IgG-HRP (Santa Cruz Biotechnology, Dallas, TX), goat anti-mouse IgG-IRDye 800 (LI-COR Biosciences) and goat anti-rabbit IgG-IRDye 680 (LI-COR Biosciences, Lincoln, NE) were commercially purchased.

### Plasmids used in this study

Plasmids and primers used in this study are listed in Additional file 1: Table S3 and S4, respectively. Plasmids were constructed using standard cloning protocols. All coding sequences of constructs used in this study were sequenced in their entirety. The plasmid pJC835 containing *HAC1^i^* gene in pRS316 was previously described [14]. The plasmids pGT0179, pGT0181, pGT0183, and pGT0185, were generated by amplifying the promoter and open reading frame of *NSG2, CUE1, SBH1*, and *EMC4* with primer pairs BN033-034, BN029-030, BN035-036, and BN031-032, respectively, from the template WT genomic DNA (gDNA). PCR products of *NSG2, SBH1*, and *EMC4* were digested with the restriction enzymes *NotI* and *NcoI* before being ligated into the corresponding restriction sites in pRS315. *CUE1* PCR product was digested with the restriction enzymes *NcoI* and *PstI* before being ligated into the corresponding restriction sites in pRS315. The plasmid pGT0288 was generated by amplifying the open reading frame of Sbh1 with primer BN027 and BN028 from WT gDNA and digested with the restriction enzyme *SfiI* before being ligated into the corresponding restriction sites in pBT3N. The plasmid pGT0350 was generated by Gibson assembly to join the promoter and open reading frame of SSS1 with primers BN013 and BN014 from WT gDNA with a 3X FLAG tag amplified with primers BN015 and BN016 from pGT0284 into pRS313. Plasmids pGT0352, pGT0445, pGT0446, and pGT0447 were generated by performing site-directed mutagenesis on pGT0183 with primer pairs BN037-BN038, PS153-PS154, PS141-142, and PS143-144, respectively, as previously described [93]. The plasmid pGT0459 was generated by sequential site-directed mutagenesis from pGT0352 using primer pairs PS143-PS144, PS141-PS142, and PS139-140 as previously described [93].

### Cycloheximide chase assay

Cycloheximide chase assay was carried out as previously described [94]. Typically, 6 OD600 units of early log phase cells were grown in synthetic media. Protein synthesis was inhibited by adding 200 μg/ml cycloheximide. Samples were taken at designated time points. Cell lysates from these samples were resolved by SDS-PAGE and transferred onto a nitrocellulose membrane. Immunoblotting was performed with appropriate primary antibodies and horseradish peroxidase-conjugated secondary antibodies or IRDye-conjugated secondary antibodies. Proteins were visualised using the ECL system (C-DiGit Chemiluminescent Western Blot Scanner) or the NIR fluorescence system (Odyssey CLx Imaging System). Values for each time point were normalised using anti-Pgk1 or anti-Tub1 as loading controls. Quantification was performed using an Odyssey infrared imaging program (LI-COR Biosciences, Lincoln, NE).

### Indirect immunofluorescence

Indirect immunofluorescence was carried out as previously described [95]. Typically, cells were grown to early log phase at 30°C in selective synthetic complete media, fixed in 3.7% formaldehyde and permeabilised. After blocking with 3% BSA, staining was performed using anti-HA (1:200), anti-LexA (1:500), anti-GFP (1:200) or anti-Kar2p primary antibody (1:1,000) followed by Alexa Fluor 488 goat anti-mouse secondary antibody (1:1,000) and goat anti-rabbit IgG-DyLight 550 (Thermo Fisher, Waltham, MA). Samples were visualised using a Zeiss LSM 710 microscope with a 100 × 1.4 NA oil Plan-Apochromat objective (Carl Zeiss MicroImaging).

### Alkaline carbonate extraction

Alkaline carbonate extraction was carried out as previously described [96]. Five OD600 units of early log phase cells were resuspended in 1.2 ml of 10 mM sodium phosphate pH 7.0, 1mM PMSF and protease inhibitor cocktail (PIC). An equal volume of 0.2 M sodium carbonate (pH 11.0) was added to cell lysates incubated 30 min at 4°C and spun down at 100,000 × *g* for 30 min, 4°C. The pellet (membrane fraction) was solubilised in 3% SDS, 100 mM Tris, pH 7.4, 3 mM DTT and incubated at 100°C for 10 min. Proteins from total cell lysate and supernatant fractions (collected from centrifuged lysate) were precipitated with 10% trichloroacetic acid (TCA) and spun down 30 min at 18,400 x g, 4°C. Proteins were resuspended in TCA resuspension buffer (100 mM T ris-HCL pH 11.0, 3% SDS).

### Proteinase K digestion assay

Fifty OD_600_ units of early log phase cells were pelleted and resuspended in 1 ml Tris Buffer (50 mM Tris pH 7.4, 50 mM NaCl, 10% glycerol, 1mM PMSF and PIC). The clarified cell lysate was spun down at 100,000 × *g* for 1 h at 4 °C. The pellet was resuspended and washed with 0.5 ml Tris Buffer without PMSF and PIC. Around ∼ 5 OD_600_ equivalent of microsomes were incubated with 1 mg/ml Proteinase K (Promega, Fitchburg, WI) and 1% Nonidet P40 substitute (Sigma-Aldrich, St. Louis, MO) when indicated and incubated at 37°C for 30 min. To quench the reaction, 5 mM PMSF was added followed by TCA precipitation. Samples were resolved by SDS-PAGE and transferred onto a nitrocellulose membrane. Immunodetection was performed with appropriate primary antibodies and IRDye-conjugated secondary antibodies. Immunoreactive species were visualised using the NIR fluorescence system (Odyssey CLx Imaging System).

### Lipid extraction and fatty acid analysis

For whole cells, 10 OD600 of early log phase cells were pelleted, washed and resuspended with ice-cold water and lyophilised using Virtis Freeze Dryer under vacuum. For lipid extraction for microsomes, 50 OD600 of early log phase cells were pelleted, washed with phosphate-buffered saline (PBS) and resuspended in 1 ml of Tris Buffer (50 mM Tris-HCL, 150 mM NaCl, 5 mM EDTA pH 8.0, 167 μM PMSF and PIC). The clarified lysate was spun down at 100,000 × *g* for 1 h at 4°C. The pellet was resuspended in 100 μl ddH_2_O and sonicated for 30 min. Lipid content was normalised to protein content using bicinchoninic acid (BCA) protein assay (Sigma-Aldrich, St. Louis, MO). Normalised microsome contents were resuspended with ice-cold ddH_2_O and lyophilised using Virtis Freeze Dryer under vacuum. Lyophilised samples were subjected to 300 μl 1.25 M HCl-MeOH (Sigma-Aldrich, St. Louis, MO) and incubated at 80°C for 1 h to hydrolyse and esterify FAs into FA methyl esters (FAME). FAMEs were extracted three times with 1 ml of hexane and separated on a gas chromatography with flame ionization detector (GC-FID; GC-2014; Shimadzu, Kyoto, Japan) equipped with an Ulbon HR-SS-10 capillary column (nitrile silicone, 25 m x 0.25 mm; Shinwa Chemical Industries, Kyoto, Japan). The temperature was held 3 min at 160°C and increase to 180°C with 1.5°C/min increments and to 220°C with 4°C/min increments.

### Fluorescence recovery after photobleaching

Fluorescence recovery after photobleaching (FRAP) was carried out as previously described [45]. Typically, early log phase cells expressing Sec63-sGFP were fixed on coverslips in Attofluor cell chambers (Thermo Fisher, Waltham, MA) with concanavalin A before rinsing thrice with ddH_2_O. Cells were imaged for 5 s followed by photobleaching a region of interest of 82 × 82 pixels at 100% intensity 488 nm laser under 5 × magnification. Subsequently, images were taken at 1.57 s intervals for a total of 160 sec. Images were acquired using a Zeiss LSM 710 microscope with a 100x 1.4 NA oil Plan-Apochromat objective (Carl Zeiss MicroImaging) with argon laser line 488 nm of optical slices 4.2 μm. ZEN black edition was used for image acquisition and analysis. Magnification, laser power, and detector gains were identical across samples. For data analysis, the fluorescence intensities of three regions of interest were measured for the duration of the experiment: the region of interest (ROI), a region outside of the cell to measure the overall background fluorescence (BG), and a non photobleached region within the cell was monitored to measure the overall photobleaching and fluorescence variation (REF). Normalised fluorescence intensity [F(*t*)*_norm_*] was calculated for each time point using Eq. 1 [97]. F(i) denotes the initial fluorescence intensities.

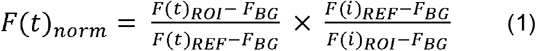

Fluorescent recovery was analysed by calculating half maximal fluorescence intensity (t_½_) using Eq. 2 [98]. F_0_ denotes the normalised initial fluorescence intensity, F_∞_ the normalised maximum fluorescence intensity and F(t) the normalised fluorescent intensity at each time point.

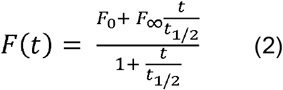

The t_½_ values were plotted using GraphPad Prism 5.0.

### Membrane yeast two-hybrid system assay

Membrane yeast two-hybrid (MYTH) assay was carried out as previously described [47]. Yeast two-hybrid screen uses the split ubiquitin two hybrid (N-terminus, N_ub_ and C-terminus, C_ub_). Briefly, MYTH bait was generated by integrating C_ub_-LexA-VP16 tag at the N-terminus of Sbh1 under the control of the promoter *CYC1* and transformed into the NMY51 yeast strain. Sbh1 tagged protein localization was verified by indirect immunofluorescence using anti-LexA antibodies against the tag described above. Seven micrograms of N_ub_G-X cDNA prey library (Dualsystems) was transformed in 35 OD_600_ units of *SBH1* reporter cells. Interactors were isolated on selective complete (SC) media lacking tryptophan, leucine, adenine and histidine complemented with 80 μg/mL X-Gal and 5 mM 3-Amino-1,2,4-triazole (3-AT) and grown for two days at 30°C. The histidine inhibitor 3-AT was used to reduce false positive colonies. Only colonies which display robust growth on selective media and a blue colour were selected for further analysis. The prey cDNA plasmids were isolated and sequenced. The list of interactors was verified via the bait dependency test, wherein all identified interactors are retransformed back into the original bait strain, together with a negative control using the single-pass transmembrane domain of human T-cell surface glycoprotein CD4 tagged to C_ub_-LexA-VP16 MYTH [49]. Interactors that activate the reporter system in yeast carrying the negative control bait were removed from the list of interactors. Yeast that harbour the prey and the bait-of-interest and did not grow were likewise removed from the list of interactors.

### Co-immunoprecipitation

Native lysis protocol was carried out as previously described [99]. Briefly, 40 OD_600_ units of exponentially growing early log phase cells were harvested and resuspended in 1 ml native lysis buffer (50 mM Tris, pH 7.5, 150 mM NaCl, 5 mM EDTA, 1 mM PIC and 1 mM PMSF). Microsomes were spun down from the clear lysates at 200,000 X *g* for 30 min, 4°C. The pellet was solubilised in native lysis buffer with 1% digitonin (Calbiochem) overnight at 4°C. The resulting lysate was cleared by centrifugation at 16,000 X *g* for 10 min, 4°C prior to immunoprecipitation. Solubilised microsomes were incubated with Protein G beads and anti-HA antibodies overnight at 4°C. Beads were washed thrice lysis buffer containing 0.5% digitonin and twice with TBS. Proteins were separated using SDS-PAGE and visualised by immunoblotting as described above.

### β-galactosidase reporter assay

The β-galactosidase reporter assay was carried out as previously described [16]. Typically, four OD_600_ units of early log phase cells were collected and resuspended in 75 μl LacZ buffer (125 mM sodium phosphate, pH 7, 10 mM KCl, 1 mM MgSO_4_, 50 mM β-mercaptoethanol). As positive control to induce the UPR, tunicamycin was added at a concentration of 2.5 μg/ml to growing WT cells 1h prior to harvest. An aliquot of 25 μl cell resuspension was transferred into 975 μl ddH_2_O and the absorbance was measured at 600 nm. To the remaining resuspension, 50 μl chloroform and 20 μl 0.1% SDS were added and vortexed vigorously for 20 sec. The reaction was started with the addition of 1.4 mg/ml ONPG (2-nitrophenyl-D-galactopyranoside; Sigma) in LacZ buffer. Then, the reaction was quenched with 500 μl of 1 M Na2CO3 when sufficient yellow colour had developed without exceeding a ten-minute reaction. The absorbance was measured at 420 and 550 nm. The β-galactosidase activity was calculated using Eq. (3).

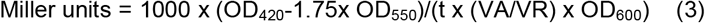

The values were then normalised to the activity of WT.

## LIST OF ABBREVIATIONS

co-IP: co-immunoprecipitation
CytoQC: cytosolic protein quality control
DMPE: N-dimethyl phosphatidylethanolamine
ER: endoplasmic reticulum
ERAD: endoplasmic reticulum-associated degradation
FA: fatty acid
LP: lipid perturbation
MMPE: N-monomethyl phosphatidylethanolamine
MYTH: membrane yeast two hybrid
NAFLD: non-alcoholic fatty liver disease
NASH: non-alcoholic steatohepatitis
PC: phosphatidylcholine
PE: phosphatidylethanolamine
SERCA: sarco/endoplasmic reticulum Ca2+-ATPase
T2D: type II diabetes
TP: transmembrane protein
UPR: unfolded protein response

## DECLARATIONS

### Ethics approval and consent to participate

Not applicable

### Consent for publication

Not applicable

### Availability of data and material

All data generated and/or analysed from this study are included in this manuscript and its additional information files

### Authors’ contributions

BSHN, PJS, and GT designed the experiments. BSHN and PJS performed the experiments with the contribution of NH, RC, and SYL. BSHN performed the experiments related to Fig. 1–3; 4a-c; 5a; S1–S3. PJS performed the experiments related to Fig. 3c-e; Fig. 4a-b; Fig. 5b-e, S2. NH performed the experiments related to Fig. 4d. RC performed the experiments related to Fig. 4a-c, SYL performed the experiments related to Fig. 4a-b. BSHN, PJS, NH, and GT contributed to the writing of the manuscript and the interpretation of the data. All authors read and approved the final manuscript.

## Acknowledgements

We are grateful to Dr. Davis Ng for providing reagents. We thank Dr. Stefan Kreft for generously providing the plasmids STK05-8-5 and STK05-5-9 [54], Yee Lin Ang and Charlie Marvalim for assisting in cloning. We thank Chengchao Xu and members of Thibault lab for critical reading of the manuscript.

## ADDITIONAL FILES

**Additional file 1:** Supplemental methods and data, Figures S1–S3, and Tables S2–S4. (PDF file)

**Additional file 2: Table S1**. List of genes upregulated transcriptionally but having lower protein abundance under LP. (XLSX file)

## SUPPLEMENTARY FIGURE LEGEND

**Figure S1.**
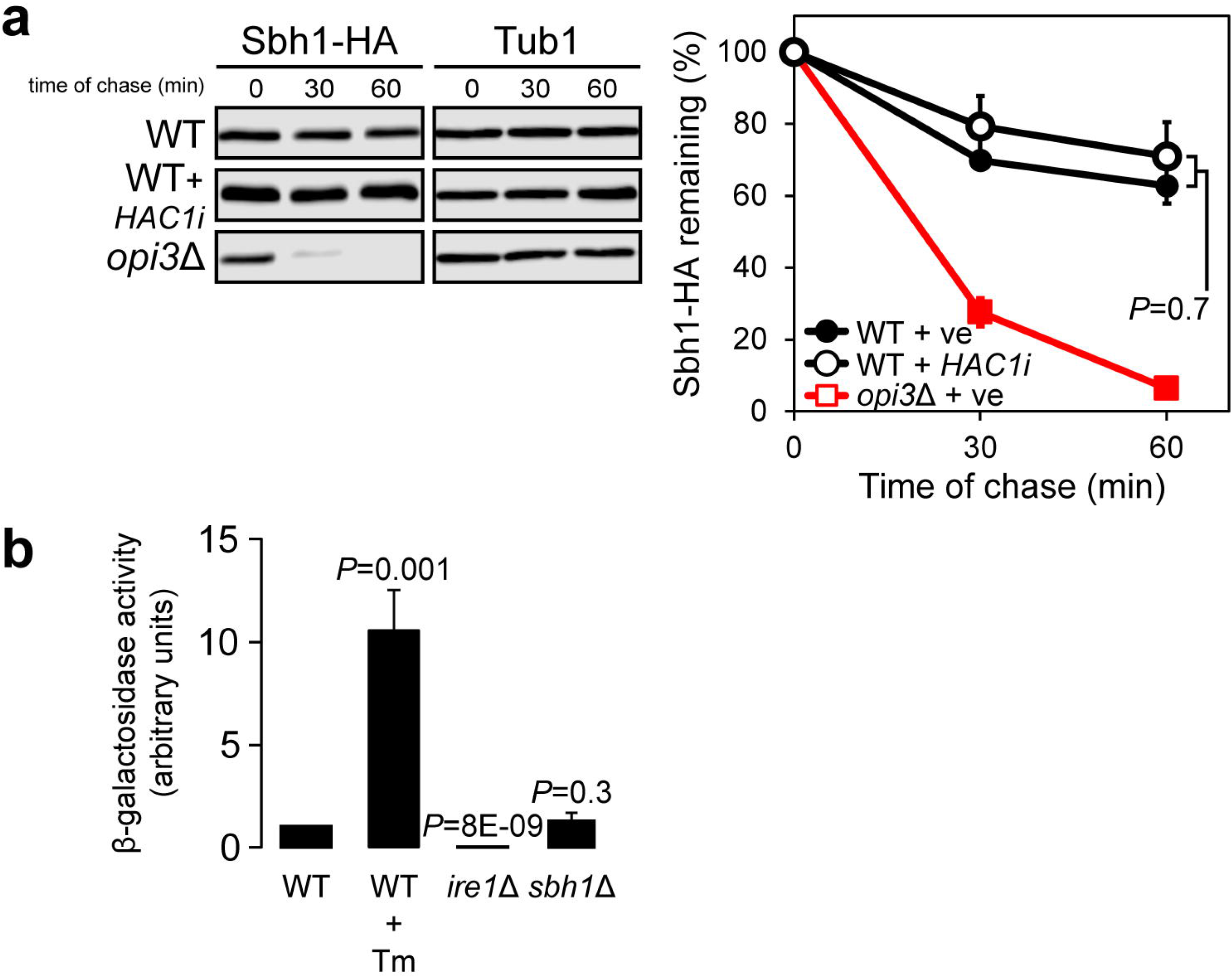
Strong activation of the UPR does not destabilise Sbh1. (**a**) The degradation of Sbh1-HA was analysed in WT and *opi3*Δ cells containing control vector (ve) or *HAC1^i^*-bearing plasmid after blocking translation with cycloheximide. Proteins were separated by SDS-PAGE and detected by immunoblotting with antibodies against the HA tag and Tub1 as loading control. (b) Cells were grown to early log phase at 30°C in selective synthetic complete media. UPR induction was measured using a *UPRE-LacZ* reporter assay. Tm, tunicamycin.

**Figure S2.**
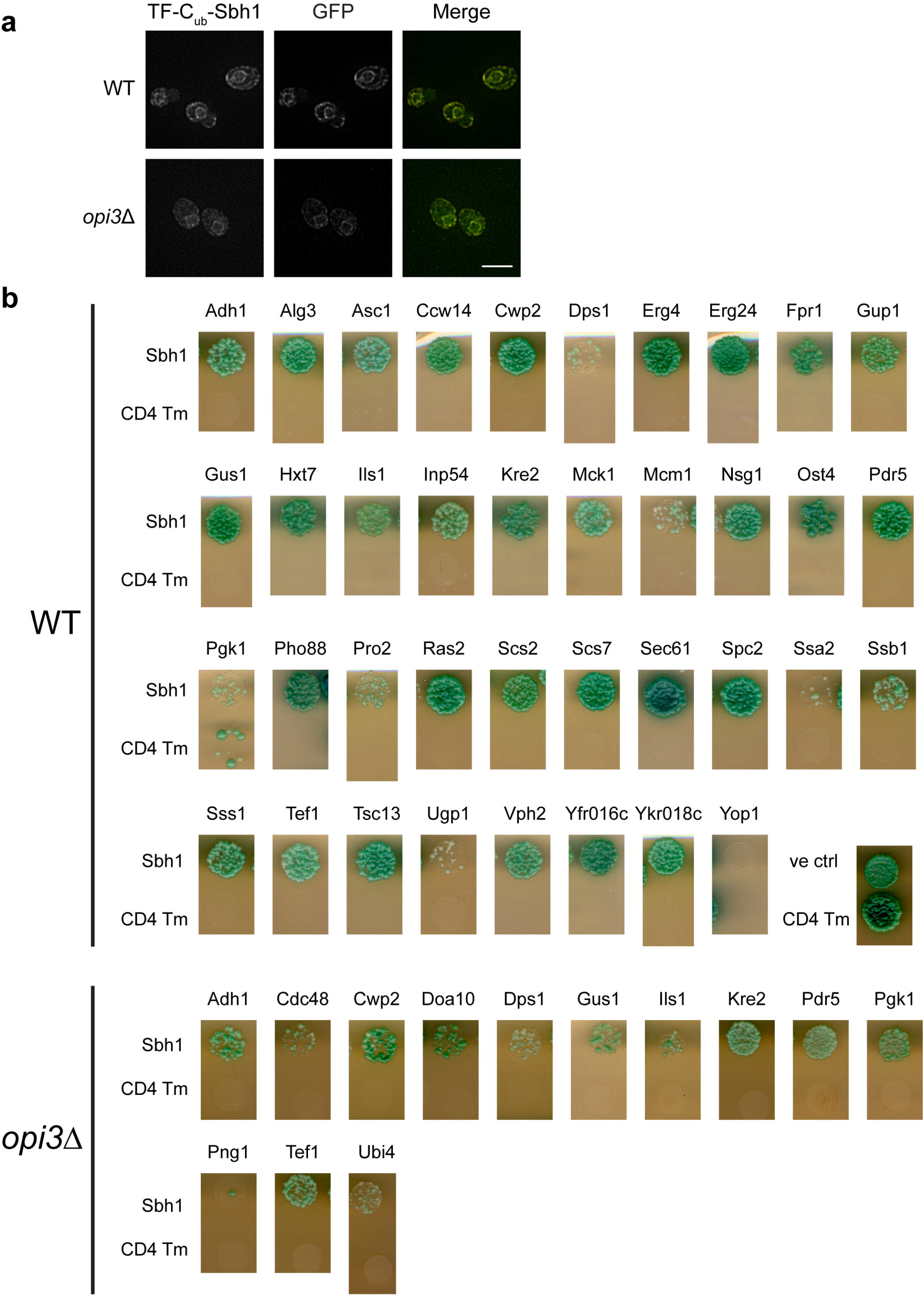
Validation of Sbh1 interacting partners. (**a**) N-termini reporter tagged Sbh1 (TF-C_ub_-Sbh1) remains localised to the ER membrane in both WT and *opi3*Δ. Protein candidates were detected using antibodies against LexA and eroGFP as ER marker. Scale bar, 5 μm. (**b**) Interacting proteins of N-tagged (TF-C_ub_-Sbh1) were retransformed with the original bait strain, together with a negative control using the single-pass transmembrane domain of human T-cell surface glycoprotein CD4 tagged to C_ub_-LexA-VP16 MYTH. Positive control of pOST1-NubI bait was used (ve ctrl). Tm, tunicamycin.

**Figure S3.**
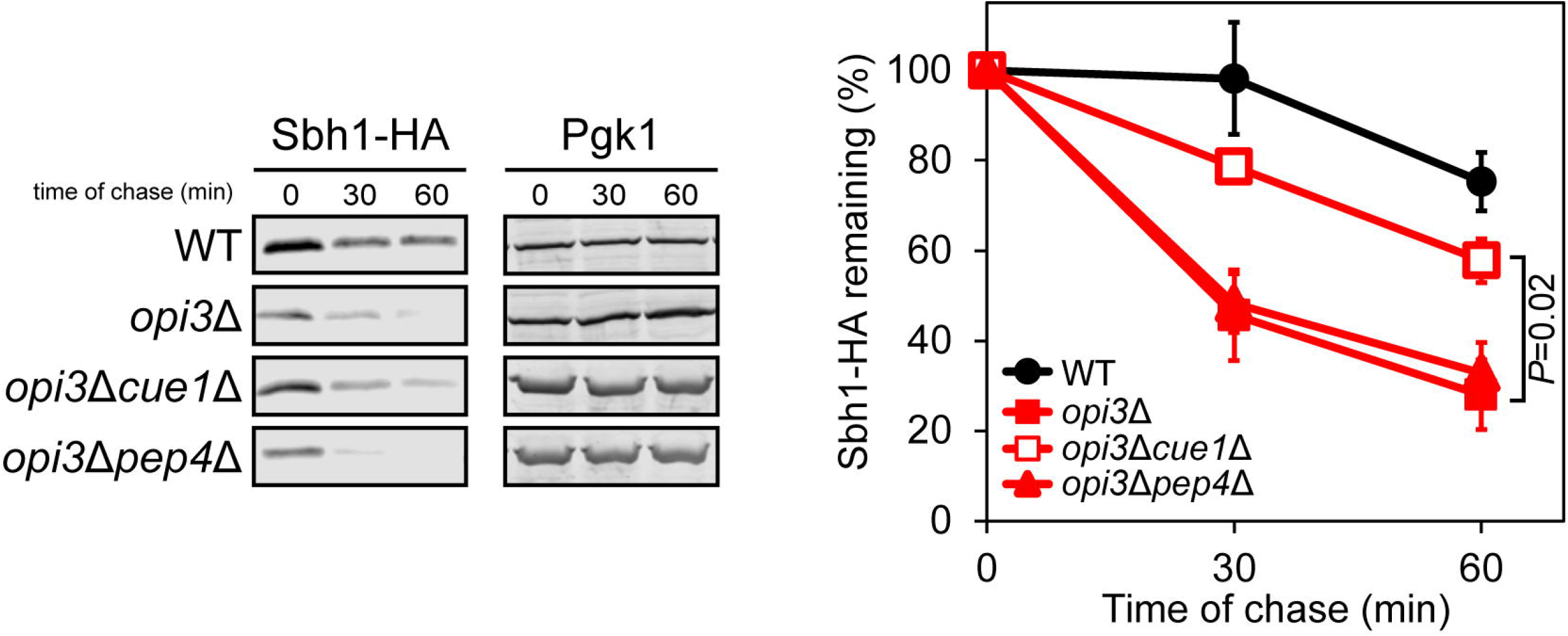
Sbh1 is degraded by the ERAD and not the vacuolar pathways. The degradation of Sbh1-HA was analysed in WT, *opi3*Δ, *opi3*Δ*cue1*Δ, and *opi3*Δ*pep4*Δ cells after blocking translation with cycloheximide. Proteins were separated by SDS-PAGE and detected by immunoblotting with antibodies against the HA tag and PGK1 as loading control.

**Table S2.** Strains used in the study

| **Strains** | **Genotype** | **Source** |
| --- | --- | --- |
| W303a | *MATa, leu2-3,112, his3-11, trp1-1, ura3-1, can1-100, ade2-1* | [14] |
| GTY68 | *MATa, opi3::KANMX, W303 background* | [13] |
| YGT0315 | *MATa, pGT0181, W303 background* | This study |
| YGT0317 | *MATa, opi3::KANMX, pGT0181, W303 background* | This study |
| YGT0318 | *MATa, pGT0182, W303 background* | This study |
| YGT0320 | *MATa, opi3::KANMX, pGT0182, W303 background* | This study |
| YGT0321 | *MATa, pGT0179, W303 background* | This study |
| YGT0323 | *MATa, opi3::KANMX, pGT0179, W303 background* | This study |
| YGT0327 | *MATa, pGT0185, W303 background* | This study |
| YGT0329 | *MATa, opi3::KANMX, pGT0185, W303 background* | This study |
| YGT0330 | *MATa, pGT0315, W303 background* | This study |
| YGT0332 | *MATa, opi3::KANMX, pGT0315, W303 background* | This study |
| YGT0374 | *MATa, pGT0183, W303 background* | This study |
| YGT0375 | *MATa, opi3::KANMX, pGT0183, W303 background* | This study |
| YGT0432 | *MATa, pJC835, W303 background* | This study |
| YGT0540 | *MATa, pGT0288, NMY51 background (his3Δ200, trp-901, leu2-3,112, ade2, LYS::(*lexAop*)4-HIS3, ura3::(*lexAop*)8-LACZ, (*lexAop*)8-ADE2, GAL4)* | This study |
| YGT0541 | *MATa, opi3::KANMX, pGT0183, NMY51 background* | This study |
| YGT0574 | *MATa, doa10::KANMX, opi3::KANMX, pGT0183, W303 background* | This study |
| YGT0575 | *MATa, hrd1::KANMX, opi3::KANMX, pGT0183, W303 background* | This study |
| YGT0576 | *MATa, usa11::KANMX, opi3::KANMX, pGT0183, W303 background* | This study |
| YGT0671 | *MATa, pGT0352, W303 background* | This study |
| YGT0672 | *MATa, opi3::KANMX, pGT0352, W303 background* | This study |
| YGT0673 | *MATa, pGT0183, pRS313, W303 background* | This study |
| YGT0674 | *MATa, opi3::KANMX, pGT0183, pRS313, W303 background* | This study |
| YGT0675 | *MATa, pGT0183, pGT0349, W303 background* | This study |
| YGT0676 | *MATa, opi3::KANMX, pGT0183, pGT0349, W303 background* | This study |
| YGT0690 | *MATa, pGT0180, W303 background* | This study |
| YGT0691 | *MATa, opi3::KANMX, pGT0180, W303 background* | This study |
| YGT0721 | *MATa, pGT0350, pGT0183, W303 background* | This study |
| YGT0722 | *MATa, opi3::KANMX, pGT0350, pGT0183, W303 background* | This study |
| YGT0725 | *MATa, pGT0350, pRS315, W303 background* | This study |
| YGT0726 | *MATa, opi3::KANMX, pGT0350, pRS315, W303 background* | This study |
| YGT0769 | *MATa, pGT0366, W303 background* | This study |
| YGT0770 | *MATa, opi3::KANMX, pGT0366, W303 background* | This study |
| YGT0771 | *MATa, pGT0368, W303 background* | This study |
| YGT0772 | *MATa, opi3::KANMX, pGT0368, W303 background* | This study |
| YGT0773 | *MATa, pGT0365, W303 background* | This study |
| YGT0774 | *MATa, opi3::KANMX, pGT0365, W303 background* | This study |
| YGT0874 | *MATa, pPS1622, pRS313, W303 background* | This study |
| YGT0875 | *MATa, opi3::KANMX, pPS1622, pRS313, W303 background* | This study |
| YGT0876 | *MATa, pGT0349, pPS1622, W303 background* | This study |
| YGT0877 | *MATa, opi3::KANMX, pGT0349, pPS1622, W303 background* | This study |
| YGT1122 | *MATa, pGT0445, W303 background* | This study |
| YGT1123 | *MATa, opi3::KANMX, pGT0445, W303 background* | This study |
| YGT1124 | *MATa, pGT0446, W303 background* | This study |
| YGT1125 | *MATa, opi3::KANMX, pGT0446, W303 background* | This study |
| YGT1126 | *MATa, pGT0447, W303 background* | This study |
| YGT1127 | *MATa, opi3::KANMX, pGT0447, W303 background* | This study |
| YGT1148 | *MATa, pGT0459, W303 background* | This study |
| YGT1149 | *MATa, opi3::KANMX, pGT0459, W303 background* | This study |
| YGT1167 | *MATa, STK05-5-9, W303 background* | This study |
| YGT1168 | *MATa, opi3::KANMX, STK05-5-9, W303 background* | This study |
| YGT1169 | *MATa, STK05-8-5, W303 background* | This study |
| YGT1170 | *MATa, opi3::KANMX, STK05-8-5, W303 background* | This study |

**Table S3.** Plasmids used in the study

| Plasmid | Encoded protein | Promoter | Vector | Source |
| --- | --- | --- | --- | --- |
| pJC31 | β-galactosidase | *UPRC-CYC1* | pRS315 | [100] |
| pPS1622 | Sec63-sGFP | *SEC63* | pRS316 | [101] |
| pJC835 | Hac1 | *HAC1* | pRS313 | [14] |
| pGT0284 | IRE1-3X FLAG | *IRE1* | pRS426 | [102] |
| pPM28 | eroGFP | *GAP* | pRS316 | [103] |
| STK05-5-9 | HA-Sbh2(S61P,S68A) | *MET25* | p413MET25 | [104] |
| STK05-8-5 | HA-Sbh121 | *MET25* | p413MET25 | [104] |
| pGT0179 | Nsg2-HA | *NSG2* | pRS315 | This study |
| pGT0181 | Cue1-HA | *CUE1* | pRS315 | This study |
| pGT0183 | Sbh1-HA | *SBH1* | pRS315 | This study |
| pGT0185 | Emc4-HA | *EMC4* | pRS315 | This study |
| pGT0288 | C<sub>ub</sub>-LexA-VP16-Sbh1 | *CYC1* | pBT3-N | This study |
| pGT0352 | Sbh1(K41A)-HA | *SBH1* | pRS315 | This study |
| pGT0350 | Sss1-3XFlag | *SSS1* | pRS313 | This study |
| pGT0445 | Sbh1(K15/17A)-HA | *SBH1* | pRS315 | This study |
| pGT0446 | Sbh1(K23A)-HA | *SBH1* | pRS315 | This study |
| pGT0447 | Sbh1(K30/31)-HA | *SBH1* | pRS315 | This study |
| pGT0459 | Sbh1(6KA)-HA | *SBH1* | pRS315 | This study |

**Table S4.**
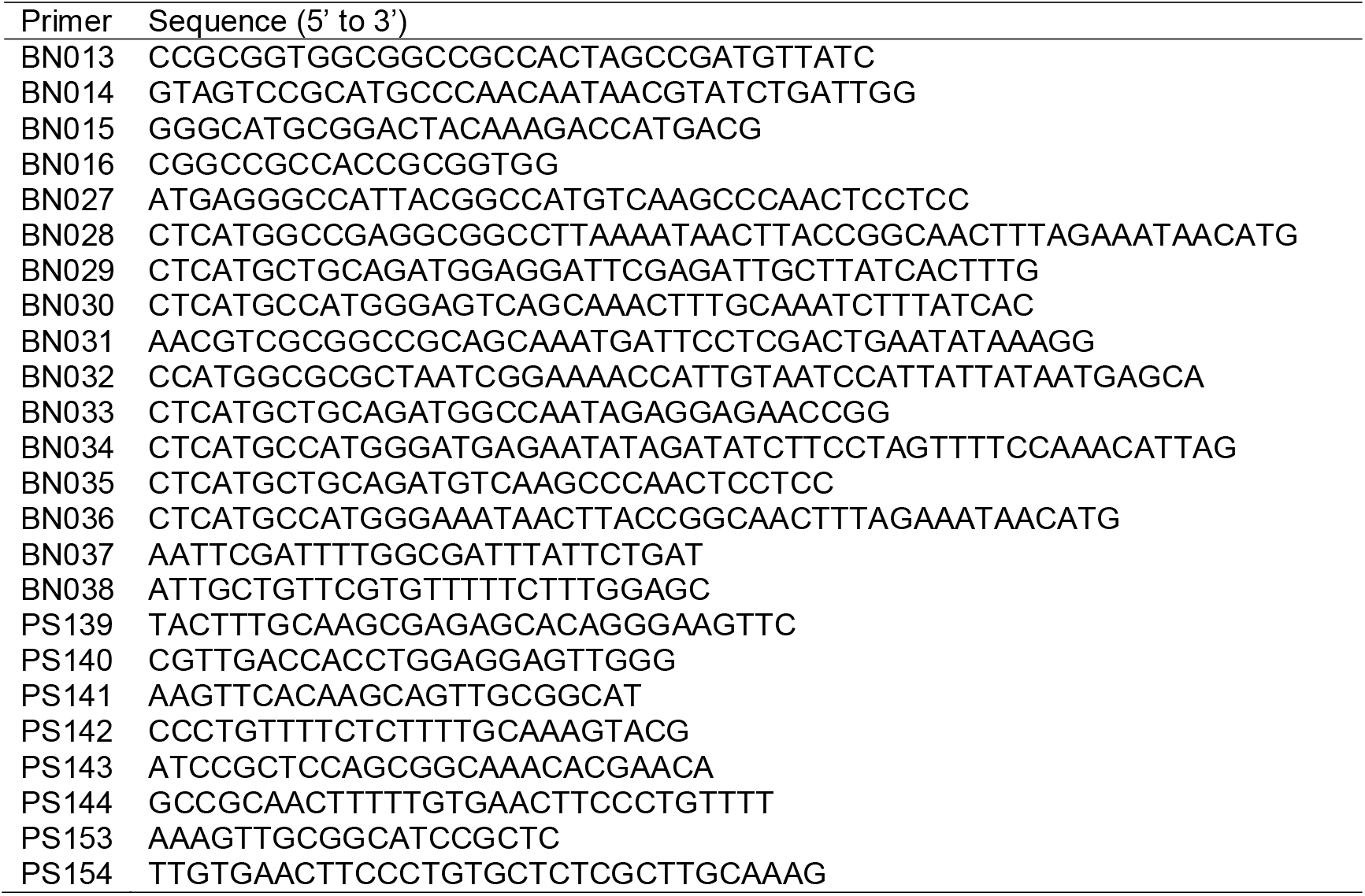
Oligonucleotide primers used in the study

